# Comprehensive 16S rRNA gene sequencing and meta-transcriptomic analyses in female reproductive tract microbiota: Two molecular profiles with different messages

**DOI:** 10.1101/2025.03.18.643876

**Authors:** Alberto Sola-Leyva, Inmaculada Pérez-Prieto, Analuce Canha-Gouveia, Eduardo Salas-Espejo, Nerea M. Molina, Eva Vargas, Apostol Apostolov, Amruta D.S. Pathare, Sergio Vela-Moreno, Susana Ruiz-Durán, Bárbara Romero, Rocío Sánchez, José Antonio Castilla-Alcalá, Merli Saare, Ganesh Acharya, Andres Salumets, Signe Altmäe

## Abstract

In recent years, high-throughput sequencing technologies have revolutionised reproductive microbiome research. Our understanding of the endometrial microbiome is primarily based on DNA-based 16S rRNA gene profiling, but DNA detection does not imply alive microbe presence. While this method is cost-effective and widely used, it has notable limitations, including the underestimation of microbial diversity, abundance, and functionality, as well as limited species-level resolution. Meta-transcriptomic analysis, an RNA-based approach, addresses these shortcomings by capturing functional transcripts that are actively expressed in living microbes. This study aims to characterize the endometrial microbiota through a dual approach, by integrating 16S rRNA gene sequencing and meta-transcriptomic analyses. By simultaneously analysing microbial composition and gene expression within the female reproductive tract samples, we seek to provide a more comprehensive understanding of its microbiota and their functional potential. A total of 49 women, aged 27–42 years, were enrolled in the study. Vaginal swabs, endometrial brushings, and endometrial biopsy samples were collected from each participant during the mid-secretory menstrual cycle phase, 6-9 days post luteinizing hormone surge. Our findings suggest that in low-microbial-biomass environments like the endometrium, the correlation between 16S rRNA gene sequencing and meta-transcriptomics is relatively weak. This highlights the limitations of microbial analysis of low-microbial-biomass samples. Alternatively, it suggests that microbial functions and genome activity are tissue-specific and dependent on the host tissue environment. Moreover, RNA-based analysis provides higher resolution in detecting certain pathogens, even within the endometrium. In conclusion, our study underscores the need to integrate genomic and transcriptomic data in microbiota studies to better elucidate critical and health-related microbiota-host interactions.

## INTRODUCTION

The human microbiota, comprising distinct communities of microorganisms localised within specific niches throughout the human body and their associated ‘theatre of activity’, is set to become a central point of medicine ^1^. Understanding the composition and function of these microbial ecosystems has redefined our understanding of the symbiotic relationship between the microbes and their host. High-throughput sequencing technologies have revolutionised the field of microbiota research, enabling comprehensive analysis of microbial communities in various body locations.

Traditionally, microbial composition has been primarily assessed using DNA-based approaches such as 16S rRNA gene sequencing and shotgun metagenomics. Although widely used, DNA-based sequencing techniques only identify microorganisms and genes in a community (i.e., microbiome), without offering insights into microbe viability or (transcriptional) activity ^2,3^. As a complementary approach, assessment of microbial gene expression can be achieved through the implementation of meta-transcriptomics, i.e., analysing the total collection of microbial RNA transcripts in a community, for understanding the functional activity, microbial composition and potential contributions to human physiology and pathophysiology ^4^. This approach transcends the static snapshot provided by DNA-based analyses, offering a dynamic perspective on microbial activity. Meta-transcriptomic profiling has advanced our understanding of the genetic composition and transcriptional activity within the vaginal microbiome ^5^, revealing that species abundance does not necessarily correlate with transcriptional activity ^6^. Additionally, meta/transcriptomics enables the prediction of impending shifts in community composition of microbes ^6^. However, technical challenges such as the short half-life of RNA ^7^, samples with high proportion of host-derived RNA compared to scarce microbial RNA, the abundance of rRNA ^4^, and the absence of a poly(A) tail in the mRNA of most prokaryotes ^8^, limits its use to certain experimental setups.

The endometrium, once believed to be a sterile environment, is now recognised as a potential niche for microbial colonization and has been associated with various gynaecological diseases ^9,10^. The modulation of the endometrial microbiome seems to be a promising clinical approach to alleviate several gynaecological conditions, such as chronic endometritis, endometriosis and risk for gynaecological cancers ^9,11,12^. Despite significant efforts to identify the core endometrial microbiota, several factors including methodological and individual-specific variations ^9,13^ have made the task difficult. To the best of our knowledge, only two studies have so far attempted to unravel the activity of endometrial microorganisms by studying their RNA content ^14,15^. Our previous RNA-based study of microbiota mapping of endometrium from healthy women identified over 5,000 microbial transcripts (bacteria, viruses, fungi, and archaea) ^15^. However, that first attempt to profile the endometrial microbiota using meta-transcriptomics failed in revealing a body-site-relevant microbial composition, mainly due to the low microbial presence. Further research relying on well-designed and controlled studies is needed to uncover the true potential of meta-transcriptomics in revealing the active microbial environment within the uterus.

This study aims to elucidate the endometrial microbiota through a dual approach combining 16S rRNA metagenomic and meta-transcriptomic analyses of the samples obtained from the same women. By simultaneously deciphering the genetic composition and gene expression of the samples from female reproductive tract, we aim to provide a more comprehensive understanding of its microbial composition and its potential functional consequences.

## RESULTS

### Non-mirroring female reproductive tract microbiome/microbiota composition using 16S metagenomic and meta-transcriptomic analyses

A total of 44 reproductive-aged women (age=34.8±3.5 years; BMI=24.2±4.2) were included into the initial study cohort (**Supplementary Table 1**). After determining the taxonomic composition across the samples by Kraken2/Bracken, *in-silico* decontamination using Decontam revealed two contaminants out of 142 (1,41%) bacterial taxa in the vaginal microbiome, six out of 554 (1,08%) in endometrial brush samples, and 10 out of 227 (4,41%) in endometrial biopsy samples (**Supplementary Table 2**).

The microbial relative abundances at genus level for both 16S rRNA gene sequencing and meta-transcriptome are shown in **Figure 1**. Overall, the vaginal microbiome was predominantly composed of *Lactobacillus*, which represents a healthy vaginal microbiome (**Figure 1A**). According to the VALENCIA algorithm, 80% of the vaginal samples were classified into *Lactobacillus*-dominant CSTs (i.e., CST I, II, III and V), while the remaining 20% had a low abundance of *Lactobacillus* spp. and were assigned to CST IV (**Supplementary Table 3**). The latter participants exhibited a distinct vaginal microbiome profile, dominated by *Gardnerella*, *Streptococcus*, *Prevotella* or *Bifidobacterium* (**Figure 1A, Supplementary Figure 1**). Similarity scores varied across CSTs, with mean values of 0.39 for CST I, 0.01 for CST II, 0.41 for CST III, 0.59 for CST IV, and 0.05 for CST V. The relatively low scores observed could be explained by the fact that most *Lactobacillus*-dominant samples were primarily composed of *Lactobacillus helveticus* (**Supplementary Figure 1**). The endometrial microbiome (DNA-based) was evaluated in endometrial brush samples (**Figure 1B**) and biopsies (**Figure 2C**). A broader microbial composition, with a marked reduction of *Lactobacillus* proportion and increase in various other bacterial genera were found in endometrial brushes and endometrial biopsies analysed by 16S sequencing, compared to vaginal samples (**Figure 1A, B and C**).

**Figure 1.**
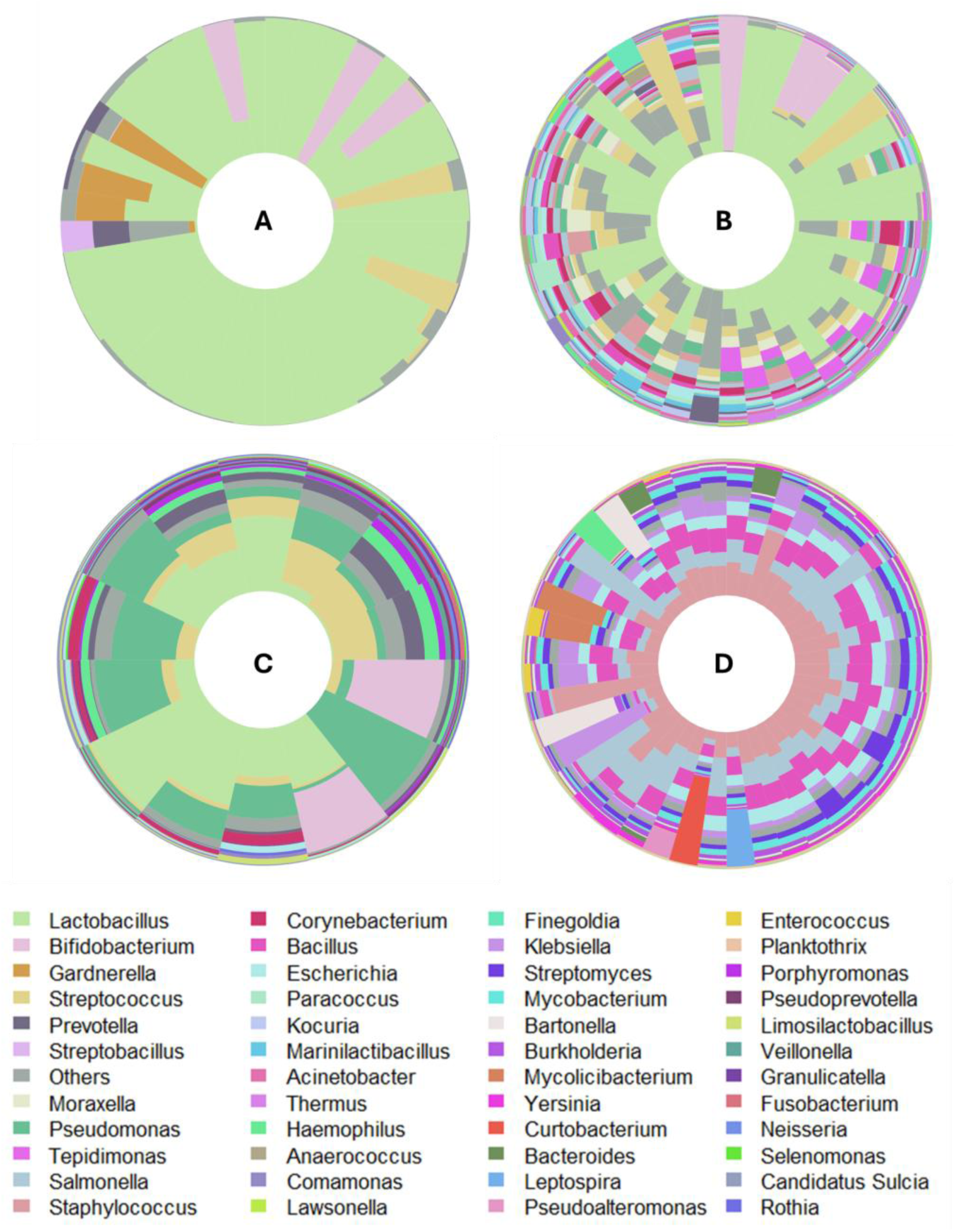
Relative abundance of bacterial genera in vagina (**A**), endometrial brush (**B**) and endometrial biopsies (**C**) detected by 16S rRNA gene sequencing (DNA-based approach). (**D**) Microbial taxa in endometrial biopsies assigned by meta-transcriptomic (RNA-based) approach. The microbial composition is plotted after decontamination. Samples on the iris plots are arranged by their rotational position around the origin of a principal component analysis (PCA) plot. Centered log ratio (CLR) transformation was applied to normalize microbial counts at the genus level. Genera with abundance less than 1% were grouped as “Others”.

**Figure 2.**
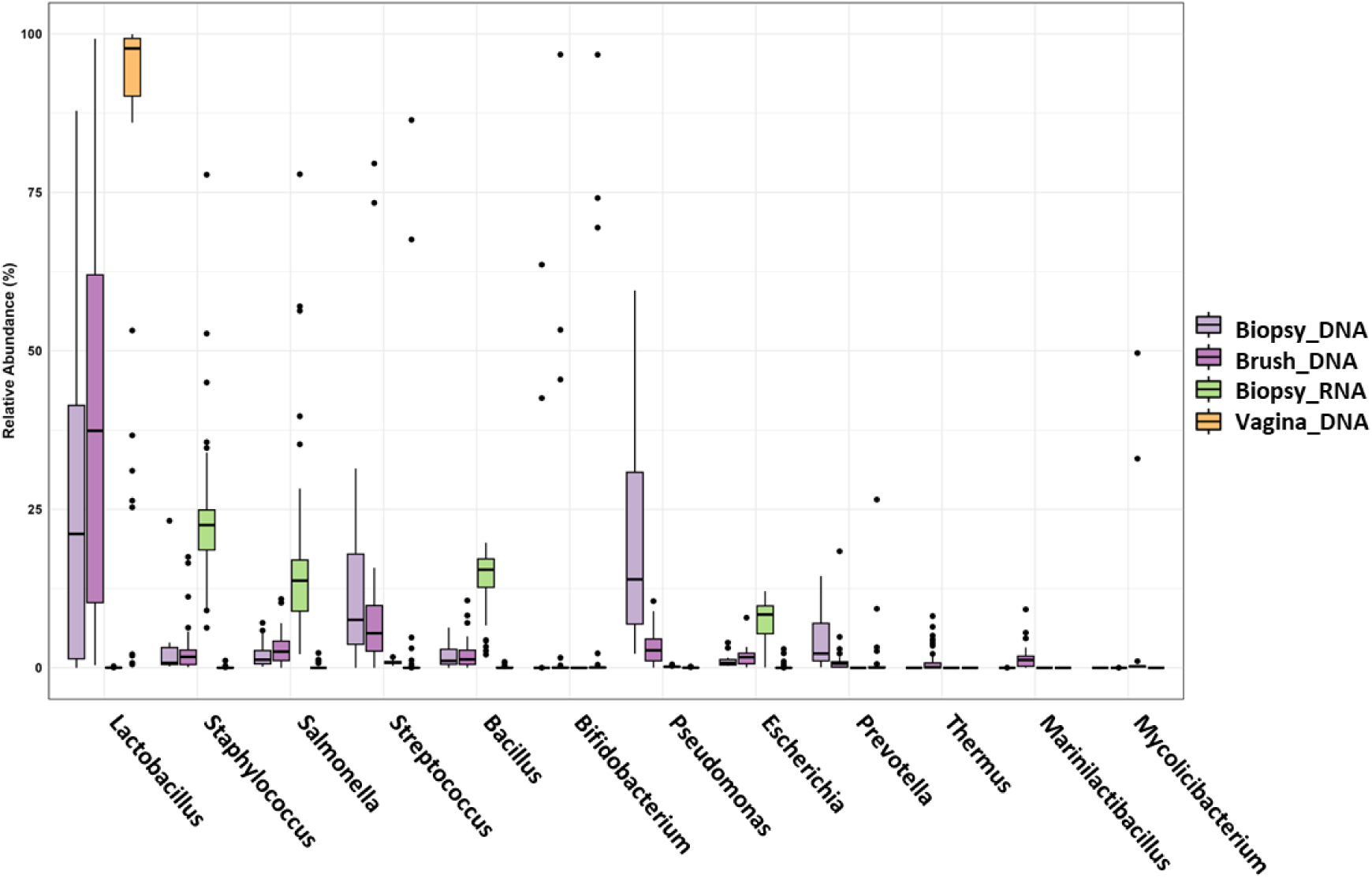
Boxplots of most abundant microbes in the vagina and endometrium detected by 16S rRNA gene sequencing and meta-transcriptomics. The boxes represent the interquartile range (IQR) of the data, with the median value indicated by the horizontal line within each box. Sample groups include 16S rRNA gene analysis of vaginal swabs (*Vagina_DNA*), endometrial biopsies (*Biopsy_DNA*), endometrial brush samples (*Brush_DNA*), and meta-transcriptomic data from endometrial biopsies (*Biopsy_RNA*). Outliers are shown as individual points. The wider spread and multiple outliers in the endometrial samples indicate substantial variability in microbial abundances across samples.

Different microbes were revealed by analysing the endometrial microbial profile by using meta-transcriptomic approach (**Figure 1D**). After decontamination, meta-transcriptomic analysis of endometrial biopsies identified 62 out of 461(13.4%) bacterial taxa as contaminants (**Supplementary Table 2**). Non-niche-specific bacterial genera like *Staphylococcus*, *Salmonella*, *Bacillus*, *Escherichia* and *Klebsiella* were identified as the most prevalent in endometrial microbial transcript analysis. This inconsistency may indicate a specific active microbial expression, with different genera being transcriptionally active within the endometrial environment or instead, but also a poor microbial assignation due to the very low number of bacterial reads compared to human reads.

To illustrate the difference in 16S rRNA gene analysis and meta-transcriptomics approaches on the identification of predominant bacteria in the vagina and endometrium, **Figure 2** displays a boxplot depicting the average relative abundance of those microorganisms that exceed 1% abundance and were shared across both anatomical sites. The relative abundance of *Lactobacillus* was higher in samples collected from the vagina compared to those from the endometrium, as indicated by the median values. Endometrial samples exhibit a wider range of variability, particularly in the meta-transcriptomic analysis of endometrial biopsies (*Biopsy_RNA* group), which showed a broad interquartile range (the height of the box) and a significant number of outliers. When comparing the microorganisms identified in endometrial biopsies, a broad range of taxa was observed depending on whether DNA or RNA was analysed. Specifically, Lactobacilli was absent in meta-transcriptome data of endometrial biopsies, indicating low average relative abundance or low genome activity in this tissue environment. On the other hand, other bacteria such as *Staphylococcus*, *Salmonella*, *Bacillus*, or *Escherichia* were more frequently detected among RNA transcripts from endometrial biopsies. Additionally, the sample collection method (brush or biopsy) significantly affected the detected microbial community. Comparing the microbiome of endometrial brushes and biopsies, we can observe that the median abundance of Lactobacilli was 12.5% higher in brushes. These results highlight that both the differential analysis of microbiome/microbiota (DNA/RNA) and the sampling method (brush/biopsy) have a significant impact on detection of microbial composition in low microbial biomass site.

### Microbial diversity analyses

Alpha diversity analyses revealed that the endometrial microbiome (both brush or biopsy samples) was significantly more diverse compared to the vagina in terms of both Shannon diversity (**Figure 3A**) and richness (**Figure 3B**) indices (all p < 0.0001). When comparing different endometrial samples (biopsy/brush), no statistically significant differences in terms of the Shannon index were found. However, endometrial brushing captured greater microbial richness compared to endometrial biopsy (**Figure 3B**; p<0.0001). Additionally, when comparing endometrial biopsy samples analysed using both 16S rRNA gene sequencing and meta-transcriptomics, both Shannon diversity and richness were significantly increased in RNA-seq analysis (all p < 0.05) (**Figure 3A and 3B).** Beta-diversity analyses on the microbial profile indicated significant dissimilarity between all the groups, i.e., vagina, endometrial brush, endometrial biopsy-DNA, and endometrial biopsy-RNA (all p < 0.001) (**Figure 3C**). The distinct clustering indicates that the sampling method (brush vs. biopsy) and the type of nucleic acid analysed (DNA vs. RNA) influence the detection of microbial community profile, which emphasizes the importance of methodological considerations in microbiome studies.

**Figure 3.**
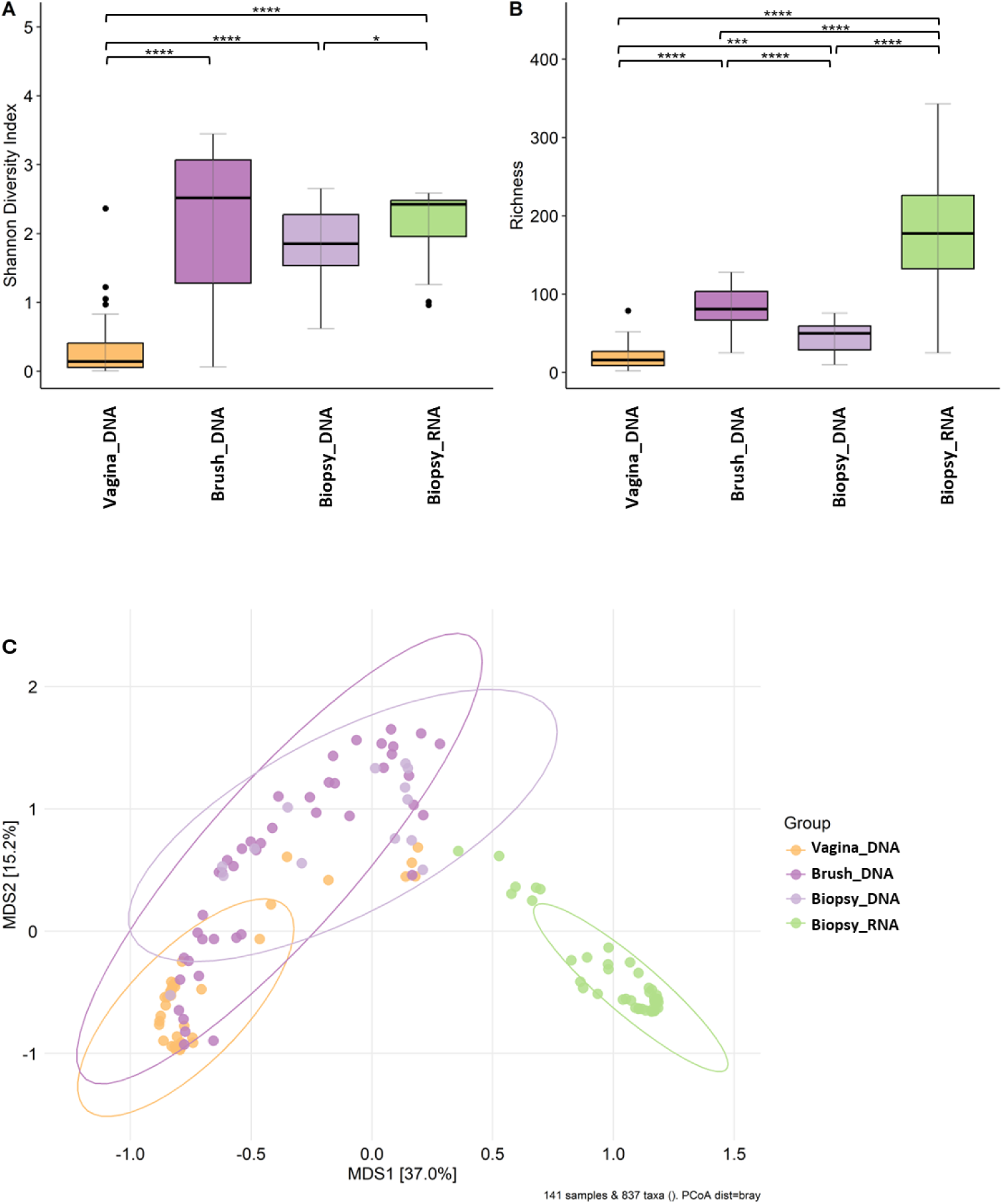
Microbial alpha and beta-diversity measures in vaginal (DNA), endometrial brush (DNA) and endometrial tissue biopsy (DNA and RNA) samples. (**A**) Shannon index and (**B**) observed richness. Significance levels: **** p < 0.0001; *** p < 0.001; ** p < 0.01; * p < 0.05; ns p ≥ 0.05. (**C**) Principal coordinates analysis (PCoA) based on the Bray–Curtis dissimilarity (Adonis PERMANOVA, all R^2^ < 0.05, all *p*-values < 0.001).

### Validation revealed the power of meta-transcriptome analysis in microbiota and dysbiosis studies of lower reproductive tract

We conducted a sub-analysis in the validation cohort where both DNA- and RNA-based techniques were applied to vaginal swabs, endometrial brush samples, and endometrial biopsies. This validation aimed to determine whether the inconsistencies observed between 16S rRNA gene sequencing (DNA) and meta-transcriptomic (RNA) taxonomic identification were due to sampling (**Figure 3C**). To assess whether performing endometrial brushing before biopsy could potentially remove active microbes, contributing to the differences observed between DNA and RNA profiles, we included a limited number of patients in the validation cohort who underwent direct endometrial biopsies without prior brushing. Notably, even in these cases, *Lactobacillus* remained undetectable at the RNA level (**Figure 4**), suggesting that its absence in endometrial biopsies was not an artifact of the sampling sequence but rather a true reflection of its low transcriptional activity in the endometrium.

**Figure 4.**
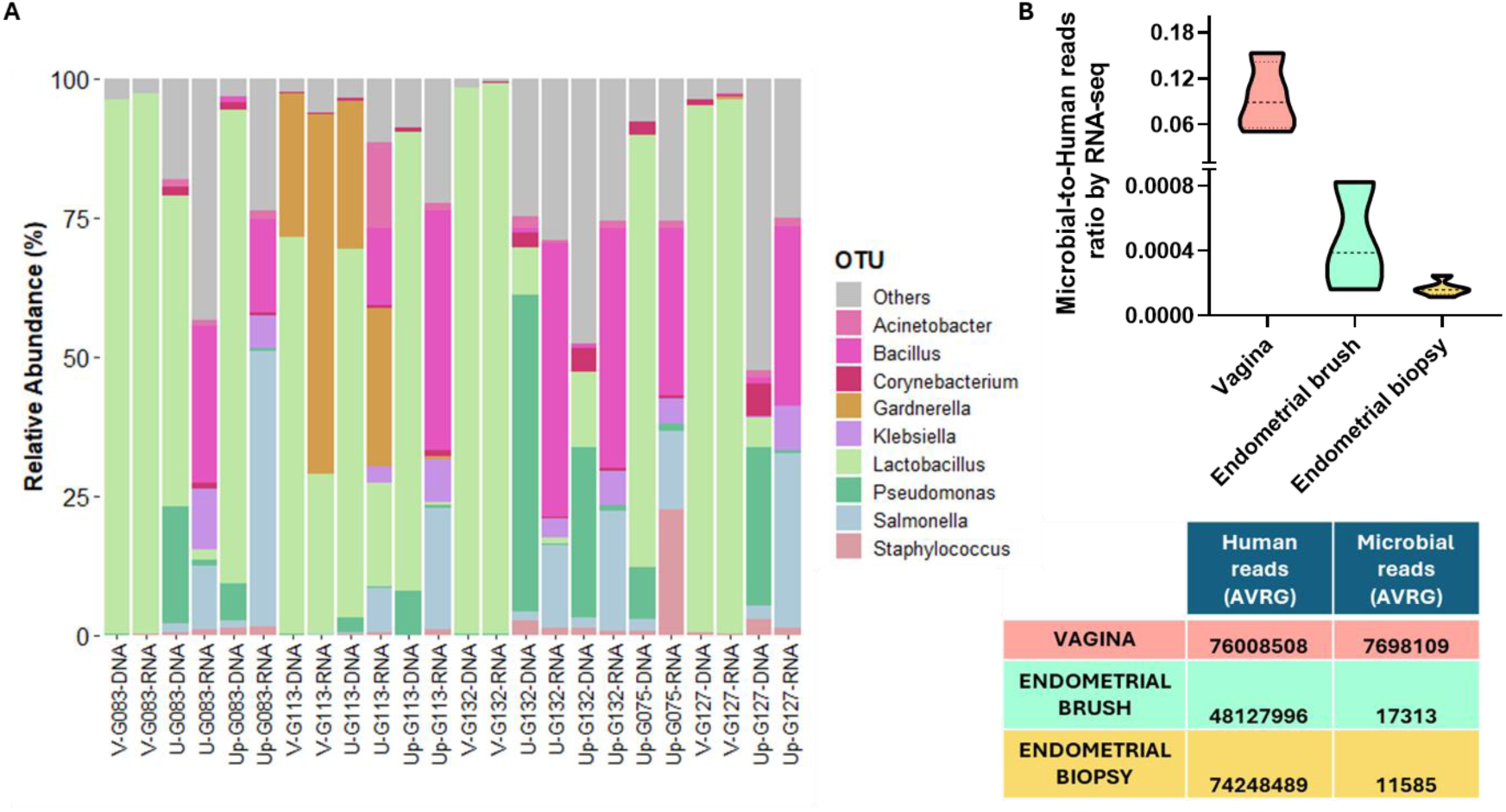
(**A**) Relative abundance of microbial composition (genus level) in the validation cohort from the vagina (V), endometrial brush samples (U), and endometrial biopsies (Up) analysed using 16S rRNA gene sequencing (DNA) and meta-transcriptomics (RNA). (**B**) Ratio of microbial-to-human reads derived from RNA sequencing (RNA-seq) in the vagina, endometrial brush, and biopsy samples. Data are presented as the mean ratio of human to microbial reads within the validation cohort. G075 and G0127 participants did not provide endometrial brush samples prior to endometrial biopsy. Vaginal swab of G075 was not able to be analysed.

**Figure 4A** illustrates the microbial relative abundances at the genus level in vaginal (V), endometrial brush (U), and endometrial biopsy (Up) samples, analysed using both DNA and RNA sequencing methods within the same samples in the validation cohort. RNA sequencing revealed a significant difference in microbial biomass, i.e., ratio of microbial-to-human reads, **Figure 4B**, with the vaginal swabs having approximately three orders of magnitude more bacterial reads than the endometrial brush and biopsy samples.

Despite differences in microbial identification between 16S rRNA gene sequencing and RNA-seq, *Lactobacillus* was consistently identified as the predominant genus in the vaginal samples, with all samples classified as *Lactobacillus*-dominant CSTs (**Supplementary Figure 2**). Notably, vaginal samples analysed by meta-transcriptomics showed a stronger alignment with VALENCIA’s CSTs, exhibiting higher similarity scores compared to those obtained by 16S rRNA sequencing (mean scores of 0.73 vs. 0.15) (**Supplementary Table 3**). Specifically, while 16S analysis predominantly identified *Lactobacillus helveticus* as the dominant species in vaginal samples, RNA-seq revealed a *Lactobacillus crispatus*-dominance in samples V-G127 and V-G132 (**Supplementary Figure 2**). Furthermore, RNA-seq provided greater resolution in detecting dysbiotic bacteria, such as *Gardnerella* and *Ureaplasma*, within the vaginal niche, compared to 16S rRNA sequencing (V-G113) (**Figure 4A, Supplementary Figure 2B**). Additionally, the meta-transcriptomic approach, but not 16S rRNA gene sequencing, was the only method capable of detecting *Gardnerella* in endometrial samples. (Up-G113, **Figure 4A**). These results demonstrate that RNA-level microbial detection provides higher taxonomic resolution and offers a better description of dysbiotic status of samples compared to 16S rRNA gene sequencing. Furthermore, the endometrial microbiota (based on RNA profile) and microbiome (based on DNA detected) were compositionally different. However, when comparing the diversity profiles in the vagina and endometrium using DNA and RNA techniques, no significant differences were observed in the validation cohort, likely due to the low sample size (**Supplementary Figure 3**).

The presence and absence analysis of the top 20 most abundant microbes in endometrium across different methodologies (16S rRNA gene sequencing and meta-transcriptomics) and sample types (endometrial brush and biopsy samples) revealed a higher number of uniquely identified microbes compared to those detected across multiple conditions (**Figure 5A**). Despite this, up to 15 shared microbial intersections were observed. Notably, *Lactobacillus* was consistently identified among the top 20 genera in all conditions, except for the meta-transcriptomic analysis of endometrial biopsies (**Figure 5B**). *Escherichia* was revealed across all methodologies and sample types. Interestingly, *Klebsiella* and *Listeria* were exclusively detected by RNA-based analysis in both brush and biopsy samples, suggesting potential transcriptional activity despite their lower DNA-based abundance. In contrast, *Bifidobacterium*, *Corynebacterium*, and *Prevotella* were solely identified at the DNA level, indicating their genomic presence without any transcriptional signals (**Figure 5B**).

**Figure 5.**
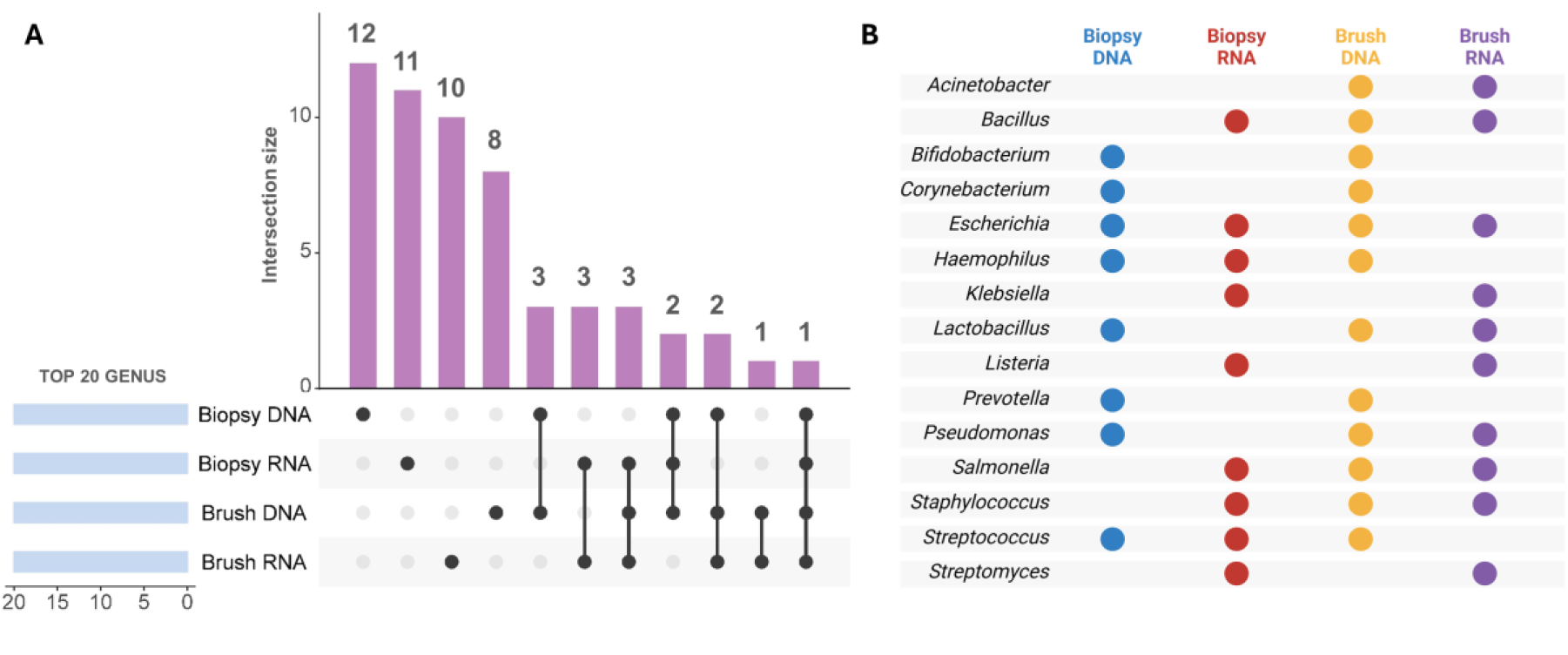
(**A**) UpSet plot illustrating the intersection of the top 20 bacterial genera (based on average abundance) detected in endometrial brush and endometrial tissue biopsy samples using 16S rRNA gene sequencing (DNA-based) and meta-transcriptomics (RNA-based). The bar plot represents the intersection size, indicating the numbers of unique and shared genera across different detection methods. Dots in the matrix represent groups included in the intersection. (**B**) Presence-absence matrix displaying the bacterial genera identified by each method, with dots indicating detection in biopsy DNA (blue), biopsy RNA (red), brush DNA (yellow), and brush RNA (purple).

The most abundant microbes in the vagina with higher relative abundance of 1%, were *Lactobacillus* and *Gardnerella* (**Figure 6A**). A comparison of the median relative abundances of *Lactobacillus* and *Gardnerella* detected using DNA- and RNA-based techniques revealed no statistically significant differences (**Supplementary Table 4A**). A total of 45 microbes were identified in vagina by using both methods both 16S rRNA gene sequencing (DNA-based) and meta-transcriptomics (RNA-based). Correlation analysis of microbes detected by DNA and RNA sequencing showed that 6 out of the 45 dually detected species exhibited a significant positive correlation (all adjusted p < 0.05) (**Figure 6B, Supplementary Table 5A**). Interestingly, the relative abundances of *Lactobacillus* or *Gardnerella* identified by DNA- and RNA-approaches positively correlated, although they did not reach statistical significance. This indicates that these microorganisms were consistently detected across DNA and RNA-based methodologies, suggesting a strong agreement between both techniques in identifying and quantifying vaginal microbial populations.

**Figure 6.**
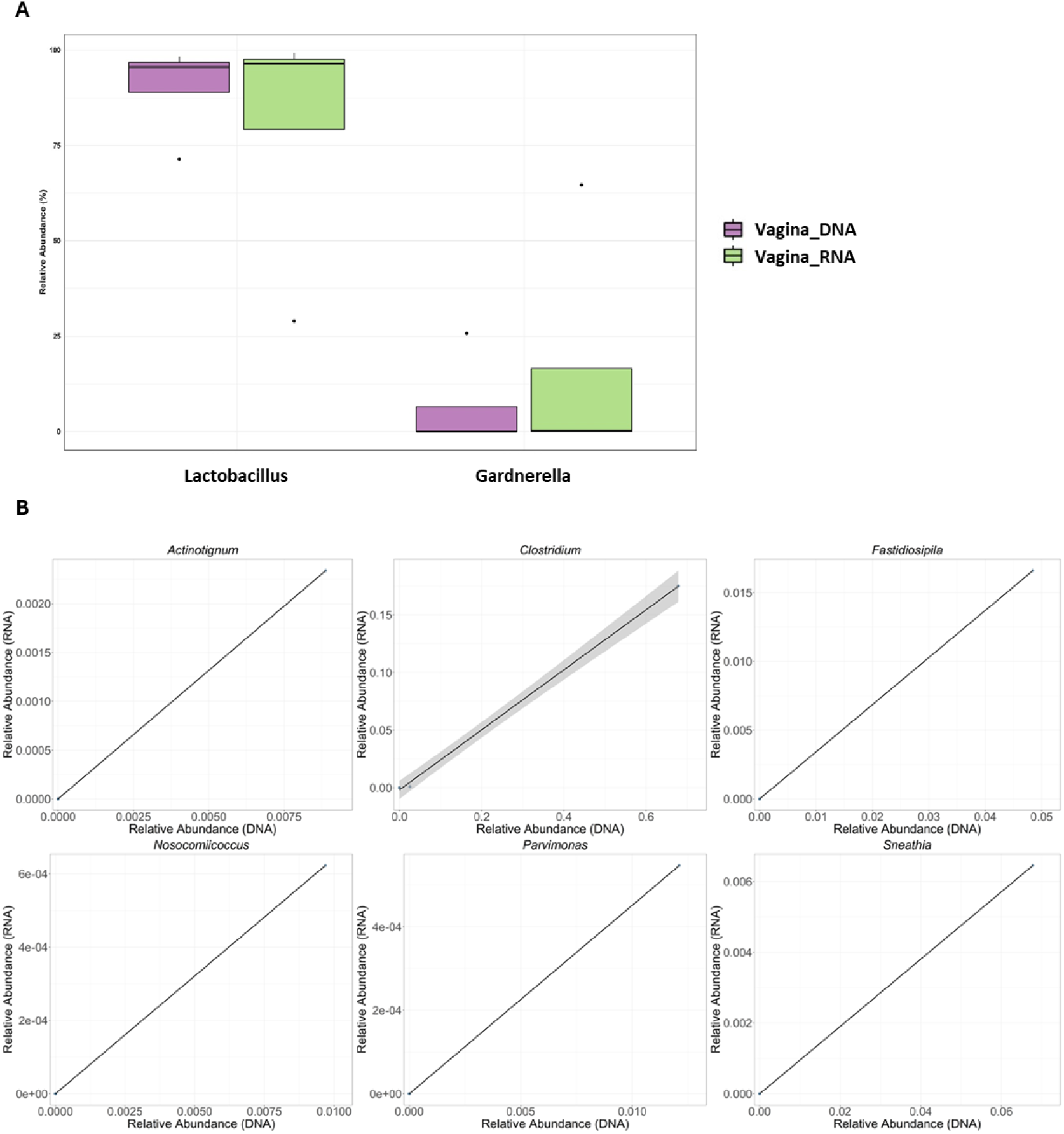
(**A**) Boxplots of relative abundance of the most abundant microbes in vaginal samples revealed by 16S rRNA gene sequencing (DNA) and meta-transcriptome (RNA) analysis. (**B**) Scatter plots of Spearman’s correlation between the taxonomic profiles of 16S rRNA gene sequencing (X axis) and meta-transcriptomics (Y axis). Significance was considered for adjusted *p*-value < 0.05 (***).

For endometrial brush and biopsy samples, the microbial landscapes revealed through the analysis of the 16S rRNA gene and meta-transcriptomic profiling were notably different (**Figure 1**, **Figure 4A**). Specifically, the median relative abundances of the most prevalent (more than 1% relative abundance) and commonly identified taxa in brush samples differed significantly between both techniques (DNA and RNA), highlighting a lack of consistency in their quantification (**Figure 7A**). Focusing on the abundance of Lactobacilli, due to their high relevance in the field, we found a lack of consistency in terms of abundance depending on the detection method used (**Figure 7A**). Despite the substantial differences, no statistically significant variation was found in the median abundance detected by DNA- vs. RNA-based techniques in endometrial brush samples, likely due to the limited sample size (**Supplementary Table 4B**).

**Figure 7.**
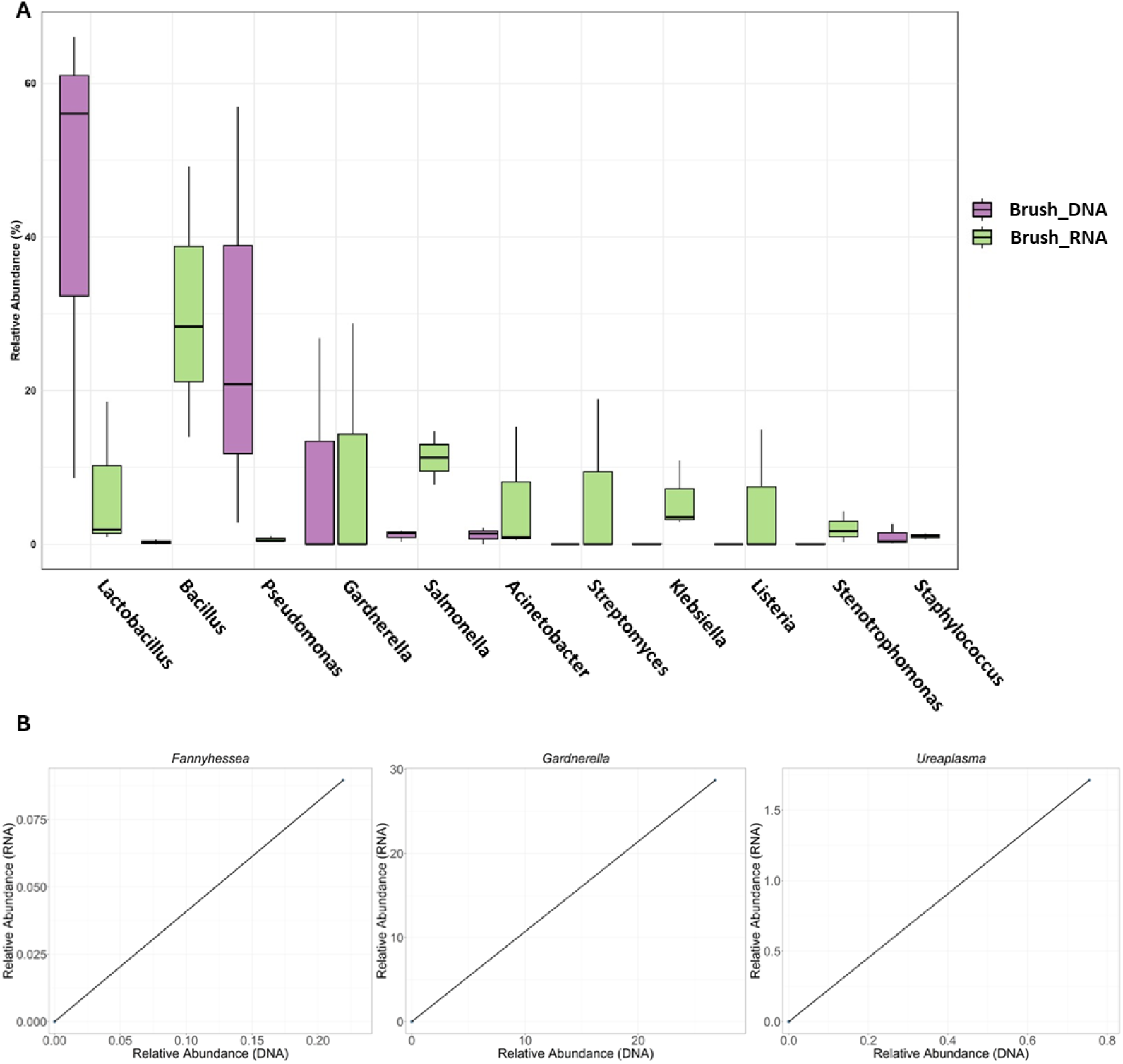
(**A**) Boxplots of relative abundance of the most abundant microbes in endometrial brush samples revealed by 16S rRNA gene sequencing and meta-transcriptome. (**B**) Scatter plots of Spearman’s correlation between the taxonomic profiles of 16S rRNA gene sequencing (X axis) and meta-transcriptomics (Y axis). Significance was considered for adjusted *p*-value < 0.05 (***).

Endometrial brush samples analysed by 16S gene/DNA sequencing and meta-transcriptomics/RNA indicated 24 microbial taxa are present in both analyses. Out of them, three bacteria were significantly and positively correlated. Specifically, *Fannyhessea*, *Gardnerella*, and *Ureaplasma* were significantly correlated by both DNA and RNA detection methods (**Figure 7B, Supplementary Table 5B)**. Specific qPCR assay performed within the validation cohort to detect pathogens, revealed that participant G113 tested positive for *Ureaplasma* together with the development of Gardnerella vaginalis and *Lactobacillus jensenii* in culture (**Supplementary Table 1**). G113 was the only participant in the meta-transcriptomic analysis of endometrial brush revealed a relative abundance of *Lactobacillus* greater than 10% (**Figure 6A**) (specifically 18% of *Lactobacillus jensenii*). This supports the results obtained by DNA and RNA sequencing methods.

The analysis of microbes in endometrial biopsies revealed the absence of *Lactobacillus* in the meta-transcriptomic data and statistically significant differences in the relative abundances of individual microbes detected by 16S rRNA gene sequencing and meta-transcriptomics. However, the abundances of *Streptococcus* and *Staphylococcus* were similar across both methods (**Figure 8A, Supplementary Table 4C**). The positive correlations observed between pathogens detected using DNA- and RNA-based methods in endometrial brush samples (**Figure 7B**) were not consistently significant in biopsy samples. Among the 18 microbes detected by both DNA and RNA techniques, only one, *Tepidimonas* remained significant. (**Supplementary Table 5C**).

**Figure 8.**
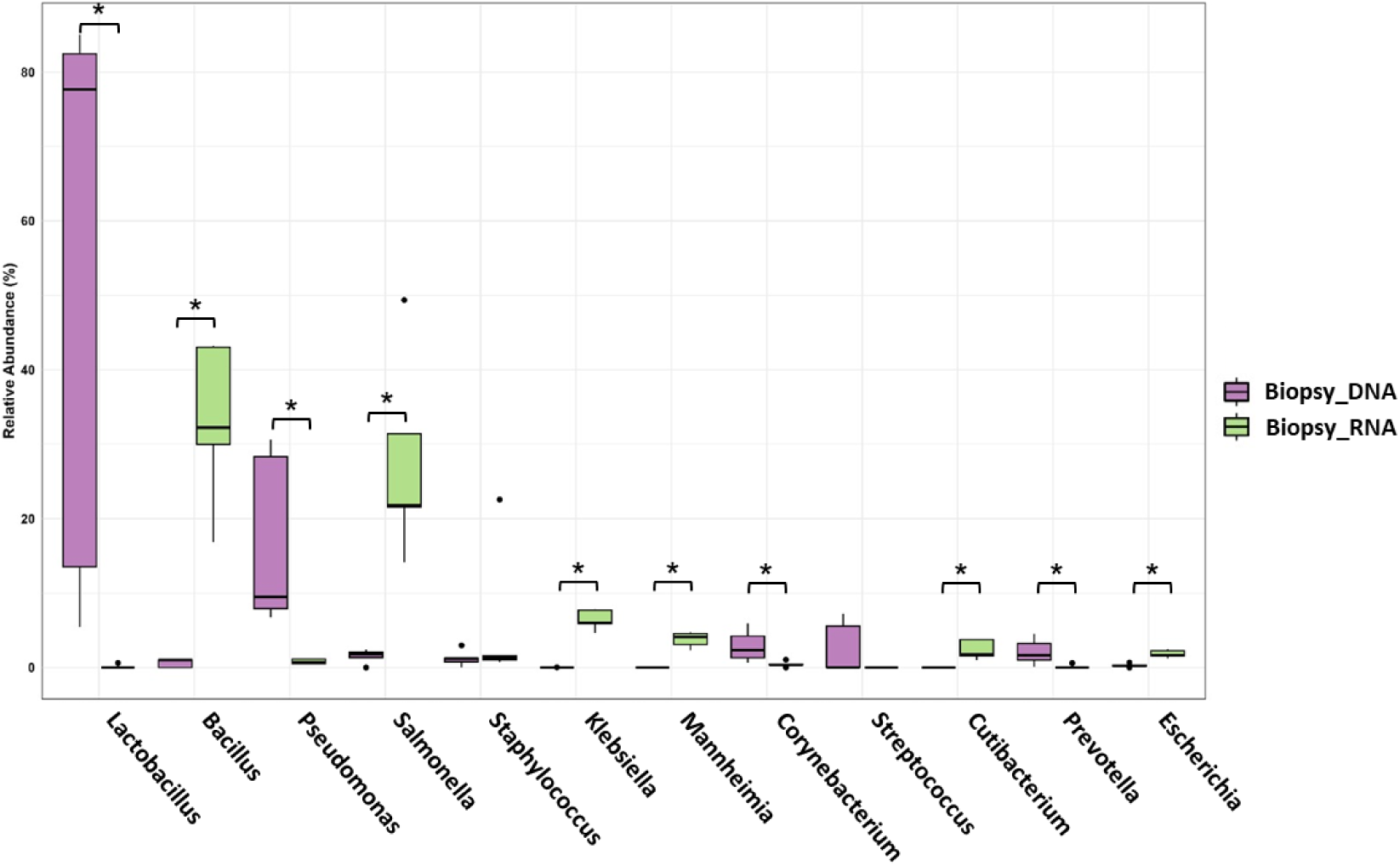
(**A**) Boxplots of relative abundance of the most abundant microbes in endometrial biopsy samples revealed by 16S rRNA gene sequencing and meta-transcriptome. Significance was considered for adjusted *p*-value < 0.05 (***).

## DISCUSSION

The endometrial microbial composition has garnered considerable attention within the field of reproductive medicine due to its potential role as a biomarker for both diagnosis and prognosis. Despite the limitations of current methodologies, characterizing microbial communities offers promising insights into reproductive health. A key focus in uterine microbiome research has been the detection of *Lactobacillus* species, whose dominance (>90%) has been linked to higher pregnancy rates ^16^. However, it remains unclear whether the bacterial DNA sequences identified by metagenomic studies correspond to live microbes. As a benchmark, the first study to map the entire active microbiota of the endometrium by identifying microbial transcripts (RNAs) described the cyclical changes in its composition throughout the menstrual cycle and identified possible host-microbiota interactions affecting endometrial receptivity ^15^. However, the absence of niche-specific microbes in meta-transcriptomic studies, combined with the lack of multi-omic approaches, makes defining an active and functional endometrial microbiota challenging. Therefore, new studies are needed to clarify whether microbes are present in the endometrium and, if so, which ones constitute the endometrial microbiota/microbiome. This study is the first to establish a continuous microbial profile from the vagina to the endometrium using a dual approach: DNA-based 16S rRNA gene sequencing and RNA-based meta-transcriptomics, validated by qPCR and bacterial culture within the same cohort and samples.

16S rRNA gene sequencing of the female reproductive tract revealed distinct microbiome in vaginal swab, endometrial brush, and endometrial biopsy samples. The method of endometrial sampling significantly impacts microbiome findings ^17^. The Tao Brush IUMC Endometrial Sampler is an endometrial sampling device featuring an outer sheath that minimizes lower genital tract contamination. Previous findings using this device identified an endometrial microbiome predominantly dominated by *Bacteroides* ^18^ and *Lactobacillus* ^19^. Investigating the endometrial microbiome from hysterectomy samples, a major surgical procedure involving the removal of the uterus revealed a distinct microbial signature in endometrial tissue, minimizing vaginocervical contamination ^20^. Opposite to the expectations, this signature was characterised by the presence of *Acinetobacter*, *Arthrobacter*, *Coprococcus*, *Methylobacterium*, *Prevotella*, *Roseburia*, *Staphylococcus*, and *Streptococcus* ^20^. In line with these findings, our results showed that *Acinetobacter* was detected at both the DNA and RNA levels only in samples collected with the Tao Brush (**Figure 5B**). Additionally, *Streptococcus* and *Staphylococcus* were transcriptionally active, as they were identified at the RNA level, suggesting biological activity beyond the mere presence of DNA sequences.

In parallel, the use of *in-silico* decontamination approaches like the Decontam R package ^21^ could be used to generate meaningful results by avoiding/reducing microbial contamination. In the studied cohort, a greater identification of contaminants was found in endometrial biopsies (4.4% of the identified bacteria) compared to approximately 1% in vaginal or endometrial brush samples. These findings underscore the necessity of integrating sampling and data analysis strategies when conducting microbiome studies in areas with potentially low microbial biomass, such as the endometrium, to minimize the influence of contaminants. Our results highlight the importance of using a sampling method that reduces the risk of contamination from the lower female reproductive tract when screening the endometrial microbial composition.

Regarding the microbiome (DNA-based) and microbiota (RNA-based) of the endometrial samples, our results consistently showed the presence of *Lactobacillus*, though with significant differences in relative abundance (**Figure 7A and 8A**). Meta-transcriptomic analysis revealed a lower representation, with *Lactobacillus* being undetectable in biopsies and present at low abundance (∼1%) in some endometrial brush samples (**Figure 1D and 5B**). These findings align with our previous study, where abundance of *Lactobacillus* species remained less than 1% in endometrial biopsies ^15^. Additionally, our prior research identified a high presence of *Klebsiella* transcripts, a result corroborated in this study, where *Klebsiella* was detected exclusively by RNA-based techniques in both endometrial brush and biopsy samples (**Figure 5B**). This bacterium has also been previously identified in the endometrium using qPCR ^22^ and 16S rRNA gene sequencing ^16,20,23^. This observation may be attributed to the brushing process potentially dislodging microbial populations in the endometrium, making them less detectable in subsequent biopsies. Despite this, some enrolled patients did not undergo endometrial brushing before the biopsy, yet they still showed no significant presence of Lactobacilli (**Figure 4A**).

The transcriptome analysis could not confirm the presence of a “core” microbiome, as identified in DNA studies. This suggests that many of the microbes detected at the DNA level may not be part of the normal endometrial microbiome and may not be transcriptionally active. A key finding of this study is the inability to detect transcriptionally active Lactobacilli in the endometrium, supporting the hypothesis that these microbes enter the uterus, but remain likely transcriptionally inactive. The presence of Lactobacilli in the human endometrium and their biological role have been subjects of research for many years ^24^, with questions raised about their ability to survive and thrive in the uterine environment, which does not provide the low pH they typically require ^25,26^, among other factors. Our results raise some doubts that Lactobacilli are not core microbes in the endometrium.

The high host RNA background and the low abundance of microbial reads in the endometrium (**Figure 4B**) complicate microbial identification ^15,27^. Specifically, our results demonstrate how profiling the microbiome using RNA-seq can reveal the vaginal microbiome with high precision. Notably, in cases of dysbiosis, RNA-seq reveals a greater pathogen activity than that identified by DNA-based techniques (**Supplementary Figure 2**). Furthermore, in eubiotic *Lactobacillus*-dominant microbiomes, the *Lactobacillus* species detected differ between both methods (**Supplementary Figures 1 and 2**). Our findings indicate that *Lactobacillus helveticus* was detected in DNA-based analyses in vagina, a species absent in the VALENCIA consensus CST database. Given that VALENCIA assigns CSTs based on species-level classification and since 16S rRNA sequencing generally does not achieve species-level resolution, the presence of *L. helveticus* in our 16S rRNA gene analyses for vaginal microbiome may reflect misclassification, since the RNA from the same samples analysed revealed instead dominance of *Lactobacillus crispatus*. Comparative genomics studies have demonstrated that *L. helveticus* and *L. crispatus* are closely related (sister) species with significant genomic overlap ^28,29^. This phylogenetic proximity complicates species-level classification using 16S rRNA gene sequencing, as their sequences share a high degree of similarity, making their differentiation unreliable with short-read sequencing approaches.

Profiling microorganisms in the endometrium remains challenging, as there is no evident correlation between the microorganisms detected in the endometrium by DNA- and RNA-based techniques. Nevertheless, several microbes were associated in cases of a dysbiotic endometrial microenvironment in endometrial brush samples (**Figure 7**), such as *Gardnerella* and *Fannyhessea* which positively correlated in endometrial brushes analysed by both DNA and RNA sequencing (**Figure 7B**). Our study revealed a greater number of vaginal bacteria whose relative abundances, determined by 16S gene sequencing and RNA transcripts, showed a strong correlation (**Figure 6**). In contrast, the number of correlations observed in the endometrial samples were then half of that found in the vagina. This disparity suggests that abundance measurements derived from DNA-based methods may not accurately reflect the true microbial activity in low microbial biomass sites, potentially leading to a misestimation of the dynamic microbial landscape within the endometrium. Alternatively, the genome activity of specific microbes is dependent on the host condition^30^, that makes the direct comparison of DNA- and RNA-based profiling approaches even more controversial.

Technically, in samples where host RNA is predominant over microbial RNA, a sizeable portion of sequencing capacity is allocated to the host. As a result, achieving adequate microbial detection requires deep sequencing, which increases both costs and computational demands ^31^. Consequently, detecting microbial sequences often requires substantially higher sequencing depth and coverage. In this study, we relied on 60–200 million reads per sample. Given that previous studies have employed sequencing depths ranging from one ^32^ to 250 million reads ^33^, our approach enhances the reliability of microbial detection by ensuring greater sequencing depth. Our previously established pipeline for microbial detection ^15^, utilizing Kraken2/Bracken, employs a k-mer-based approach. This method has demonstrated greater sensitivity in taxonomic classification of meta-transcriptomes, particularly in samples with a high human-to-bacteria cell ratio^34,35^.

The novelty of this study lies in its parallel dual analysis using DNA- and RNA-based techniques along the female reproductive tract to characterize the microbiome and microbiota. This analysis revealed that in low-microbial-biomass environments like the endometrium, the correlation between the two techniques is relatively weak, highlighting how the detected microbial landscape varies depending on the chosen approach. Still, RNA-based analysis provided higher resolution in detecting certain pathogens, even in the endometrium. Our analysis revealed that microbes related to vaginal dysbiosis were also transcriptionally active, demonstrating the significant advantage of RNA-based methodologies. Although the endometrial microbiome, primarily studied using DNA sequences, has been suggested as a prognostic biomarker for female reproductive health, future studies should more clearly differentiate between the microbiome and microbiota. However, as RNA-based technologies for low microbial biomass are still developing, larger studies are needed to confirm and support the findings of this study.

## MATERIAL AND METHODS

### Study population and design

This cross-sectional study was approved by the Ethics Committee of the Junta de Andalucía (CEIM/CEI 0463-M1-18r). All participants provided written informed consent before enrolment.

The study cohort consisted of forty-four reproductive-aged women (27 – 42 years old) undergoing assisted reproductive treatment (ART) at the Reproductive Unit at Virgen de las Nieves University Hospital, Granada, Spain, between March 2019 and April 2023 (**Figure 9**). Additionally, five women undergoing ART in the same unit were selected for a validation cohort. They provided vaginal, endometrial brush, and endometrial biopsy samples for parallel 16S rRNA gene sequencing and meta-transcriptome analyses. For two women, in the validation cohort, endometrial brush samples were not taken. One vaginal swab from the validation cohort could not be analysed. The aim of the validation was to determine whether different microbial detection techniques affect the intra-individual microbial signature, comparing in more detail the DNA and RNA-based techniques among all sample types (along the reproductive tract) which was not possible for the full cohort (**Figure 9**).

**Figure 9.**
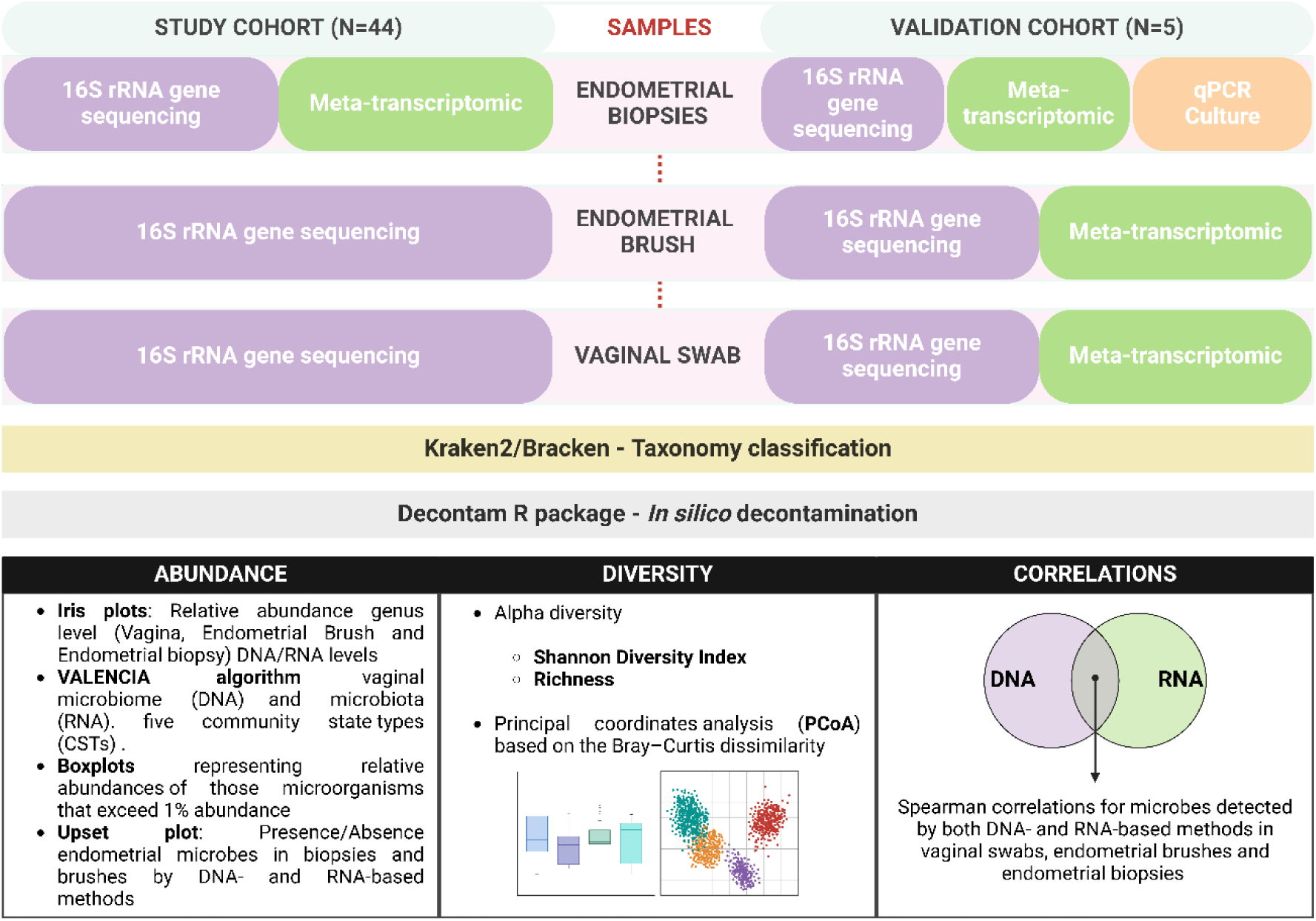
Workflow for characterizing the microbiome (16S rRNA sequencing; DNA-based) and microbiota (meta-transcriptomics; RNA-based) in vaginal swabs, endometrial brushes, and endometrial biopsies. Taxonomy was assigned using the Kraken2/Braken methodology, and *in silico* decontamination was performed with the Decontam R package. Characterization included abundance, diversity, and correlation between techniques. The study cohort consisted of 44 reproductive-age women. In the validation cohort (N=5), a dual approach (DNA- and RNA-based) was applied to all three sample types. Additional validation analyses for endometrial biopsies were performed using real-time PCR (qPCR) and microbial culture.

Infertility causes of all participants is presented in **Supplementary Table 1** and included endometriosis diagnosed by laparotomy or laparoscopy surgery; recurrent implantation failure defined as repeated implantation failure after the transfer of three good-quality embryos; unexplained infertility; single women undergoing ART; male factor infertility considering the recommendations of the World Health Organization manual for semen analysis; or tubal obstruction or damage (i.e., tubal factor infertility).

Participants had not received antibiotics within the three months preceding sample collection. Exclusion criteria comprised of patient’s age of ≥43 years, undergoing hormone therapy, having gynaecological tumors, systemic diseases, pelvic inflammatory disease, or other pelvic pathological conditions, other than endometriosis. To synchronize the collection of endometrial samples, all samples were collected at the mid-secretory phase, i.e., 6-9 days post-luteinizing hormone (LH)-surge (LH+6-9), where the ovulation day was estimated with a digital ovulation test (Clearblue, Swiss Precision Diagnostics GmbH). The study cohort women provided vaginal, endometrial brush, and endometrial biopsy samples for analyses as described in **Figure 9**.

### Sample collection

During the visit to the clinic, a vaginal swab (eNAT® 606CS01R; COPAN), endometrial brush and endometrial biopsies were collected during the mid-secretory phase of the cycle (LH surge+6-9 days) by the same gynaecologist. To ensure minimal contamination with bacteria from the lower reproductive tract, endometrial brush was performed using Tao Brush IUMC (Cook Medical) that was carefully closed within the uterine cavity after sample collection. Subsequently, the samples obtained from the brushing were stored in Copan eNAT transport system (eNAT® 606C; COPAN) and stored at a temperature of - 80°C. Subsequently, during the same procedure an endometrial biopsy was taken using curette device (Gynétics, Medical Products). The endometrial biopsies were divided into routine histology evaluation and microbiological analysis. Two additional aliquots were collected and immediately frozen in cryovials using the gas phase of liquid nitrogen for DNA and RNA extraction. These samples were then stored at a temperature of −80°C until further analysis.

### DNA and RNA isolation

Microbial DNA was extracted from 300 μL of vaginal, and endometrial brush specimens resuspended in 2ml of Copan eNAT transport medium. Approximately 50 ng of endometrial biopsy tissue were used to extract microbial DNA. Microbial DNA extractions were performed by QIAamp UCP Pathogen Mini Kit (Qiagen) and bead-beating on a TissueLyser II according to the manufacturer’s instructions and automated onto a Hamilton STAR robotic platform. DNA quality and purity (230/260/280 nm) were evaluated by NanoDrop ND-1000 Spectrophotometer (Thermo Fisher Scientific) and DNA quantification with Qubit 4 fluorometer (Thermo Fisher Scientific).

RNA from vaginal swabs and endometrial brushes was extracted with RNeasy Plus Universal Mini Kit (Qiagen). miRNeasy Micro kit (Qiagen) was used to isolate RNA from up to 30 mg of endometrial tissue followed by Ribo-Zero plus kit (Qiagen) processing for removing rRNA. RNA quality and quantity were checked on Bioanalyzer TapeStation 4200 (Agilent) with RNA ScreenTape (Agilent). Additional cleanup using the RNeasy MinElute Cleanup Kit (Qiagen) was performed on RNA from vaginal swabs due to DV200 values less than 50%.

### 16S rRNA gene sequencing

The microbiome was profiled by amplifying the V4 hypervariable region using the 515F (5’-GTGYCAGCMGCCGCGGTAA-3’) and 806R (5’-GGACTACNVGGGTWTCTAAT-3’) primers. All PCR reactions were performed in a 25 μl total volume containing 12.5 μl 2X KAPA HiFi Hotstart ready mix (KAPA, Biosystems), 5 μl of each primer (1 mM), and 2.5 μl of undiluted extracted DNA under the following cycling conditions: initial denaturation at 95°C or 3 min, followed by a cyclic 3-step stage consisting of 35 cycles of denaturation at 95°C for 30 s, annealing at 55°C for 30 s, and extension at 72°C for 5 min. The resulting amplicons were subjected to electrophoresis using 2% agarose gel, the size of each amplicon was around 380 pb, and DNA quantification was measured by Qubit 4 fluorometer (Thermo Fisher Scientific). Next, a double purification with magnetics beads was carried out (AMPure XP, Beckman Coulter). The last quality control was evaluated using HS Bioanalyzer (Agilent Technologies) to assess the size of the library and the absence of primer peaks. Illumina Nextera library preparation was performed according to the manufacturer’s specifications, combining PhiX phage (20%) with the amplicon library to give diversity to the run. The final library was paired-end sequenced (2 × 300 bp) using a MiSeq Reagent Kit v.3 on the Illumina MiSeq sequencing system as per the manufacturer’s instructions (Illumina).

### Microbiological evaluation by qPCR and bacterial culture

Microbiological analysis in the validation cohort was performed according to the routine workflow of the hospital microbiology laboratory ^36^. Briefly, using a semi-quantitative multiplex PCR assay (qPCR) *Lactobacillus* spp., *Gardnerella vaginalis*, *Enterococcus faecalis*, and a range of pathogens, including *Chlamydia trachomatis*, *Neisseria gonorrhoeae*, *Trichomonas vaginalis*, *Ureaplasma parvum*, *Ureaplasma urealyticum*, *Mycoplasma genitalium*, and *Mycoplasma hominis* were detected. Microorganisms grown in habitual cultures were identified using MALDI Biotyper (Bruker Daltonics) or MicroScan (Beckman Coulter) systems.

### RNA sequencing – meta-transcriptomics

The sequencing library was prepared using Illumina Stranded Total RNA Prep, Ligation with Ribo-Zero Plus protocol (Illumina), following manufacturer instructions. The input contained RNA normalised to 1,000 ng for endometrial samples, and between 500-1000 ng total for vaginal samples. Libraries were quantified with Qubit 4 fluorometer (Thermo Fisher Scientific), and quality control was performed using the fragment analyzer TapeStation 4200 (Agilent) with high sensitivity D1000 ScreenTape (Agilent). RNA sequencing was performed on the NovaSeq 6000 system (Illumina) using an S4 flow cell with paired-end sequencing (100 bp x 2) and a final depth of 60–100 million reads per sample. The RNA sequencing from the validation cohort was conducted on NextSeq 1000 Illumina platform with paired-end sequencing (50 bp x 2) and a final depth of 60–200 million reads per samples.

### Bioinformatics methods

After 16S rRNA or RNA sequencing, the resulting fastq files undergo quality evaluation using FastQC. A per sequence quality score of 30 (Q30) was used, which is a benchmark for quality in next-generation sequencing and highly reduces the probability of error in a read. Reads that successfully passed quality control were processed with the pipeline developed using Nextflow ^37^. In this pipeline, Kraken2 ^38^ was the aligner used to classify the reads in their corresponding taxa. To perform the taxonomic classification, the standard database (October 2023 version) provided by Kraken2 was used as a reference, setting the minimum hit groups option to 3 and using the paired flag. As a result, a Kraken report file was obtained for every sample, which contained the sample reads classification at multiple levels. To estimate the microbes’ abundance at a single level in the taxonomic tree, the Kraken reports were used as an input for Bracken ^39^. This analysis used the same database as the Kraken2 process as a reference, using a k-mer length of 50. As a default, the reads were classified to genus level and the reads threshold was set to 10, meaning that any microbial genus with 10 or less reads in the input Kraken report would not receive any additional reads from higher taxonomy levels when distributing reads for abundance estimation.

### Microbial decontamination

Given the need for stringent contamination control when characterizing low microbial biomass sites such as the upper reproductive tract, we employed an *in-silico* decontamination approach using Decontam R package (v.1.6.0) ^21,40^. This method identifies contaminating DNA features, allowing us to remove them in downstream analyses. Decontam presents two contamination identification methods: the “frequency method”, based on the DNA concentration; and the “prevalence” method, in which the presence/absence of a feature in the samples is compared to that in negative controls to identify contaminants ^21^. Recently, Decontam has been validated for meta-transcriptomics analysis, by applying a modified frequency method based on the total read counts per sample as material genetic quantity input ^41^. We set the Decontam threshold at 0.1 by default to identify contaminating phylotypes. To ensure comparability, the same frequency approach was used to decontaminate 16S rRNA data.

### Statistical analyses

Statistical analyses were performed using R statistical software v.4.2.1 under RStudio v.2022.07.2. The microbial data generated after taxonomy assignation in both, 16S rRNA gene sequencing and meta-transcriptomic data were aggregated to genus level for diversity and abundance comparisons. The normality of the variables was assessed using the Shapiro-Wilk test. The relative abundances of the identified genera did not conform to normal distribution and were therefore analysed using the nonparametric Mann-Whitney *U* test. The Benjamini-Hochberg method, the false discovery rate (FDR), was utilised to calculate adjusted *p*-values for multiple comparisons. Differences were considered statistically significant between groups when adjusted p < 0.05. Alpha-diversity indices, including the Shannon diversity index and observed genera richness, were calculated utilizing the “diversity” function from the “vegan” package in R. Differences in these diversity indices and abundances among sample groups were assessed using the Mann-Whitney U test. Additionally, alpha-diversity comparisons between different sample types from the same women were conducted using the Wilcoxon signed-rank test for paired data. For beta-diversity analysis, Bray-Curtis dissimilarity was computed with the “vegdist” function in R, and Permutational Analysis of Variance (PERMANOVA) was conducted using the “adonis” function from the “vegan” package to evaluate differences in microbial community composition. Further, the vaginal microbiome was classified into the five community state types (CSTs) established by the VALENCIA algorithm ^42^. VALENCIA is a well-established, nearest-centroid based method that assigns vaginal samples to five CSTs based on their similarity score to reference centroids. Species from *Lactobacillus*, *Gardnerella*, *Prevotella*, *Atopobium* and *Sneathia* genera were identified for proper CST assignment. For validation samples, a Spearman correlation test was also performed between relative abundances for each pair of genera from the DNA and RNA dataset and was represented in scatter plots. The Spearman correlation values indicated the degree to which microbial DNA relative abundance predicts RNA abundance for a given genera pair.

## DATA AVAILABILITY

Additional data of the study is included in the Supplementary Materials. Further inquiries can be made to the corresponding authors to get access to the background data.

## AUTHOR CONTRIBUTIONS

**A.S.L.** participated in the conceptualization of the project, served as the project manager, organized the sample collection, standardized the laboratory protocols (DNA/RNA extraction, library preparation, and sequencing), participated in the analysis of the samples, and wrote the manuscript. **I.P.P.** participated in the sample collection, optimized the data analysis, and implemented figure creation. **A.C.C.** contributed to the sample collection, data analysis, and visualization. **E.S.E.** optimized the data analysis and implemented figure creation. **N.M.M.** and **E.V.** contributed to the sample collection. **A.A.**, **A.D.S.P.**, and **M.S.** contributed to the validation cohort sequencing. **S.V.M.** optimized the data analysis. **S.R.D.**, **B.R.**, **R.S.**, and **J.C.A.** provided clinical guidance and contributed to sample collection. **G.A.** and **A.S.** contributed to manuscript writing, provided resources, and acquired funding. **S.A.** was responsible for conceptualization, obtaining ethical permission, essential resources, supervising the project, and revising the manuscript. All authors discussed the results and approved the final version.

## ACKNOWLEDGMENT

We sincerely thank the clinicians and research nurses for their invaluable support in patient recruitment and biopsy collection. We also extend our deepest gratitude to the patients for generously providing the essential samples that made this research possible. We are also grateful to Fernando Barba-Pareja, Luis Javier Martinez-Gonzalez, Esperanza Santiago and Juan Miguel Guerrero from GENYO (Pfizer-University of Granada-Junta de Andalucía Centre for Genomics and Oncological Research) for their excellent collaboration. We acknowledge the research support by Copan Italia S.p.A Inc., and Clearblue, SPD Swiss Precision Diagnostics GmbH.

