## Supplementary Figure for "Comprehensive 16S rRNA gene sequencing and meta-transcriptomic analyses in female reproductive tract microbiota: Two molecular profiles with different messages"

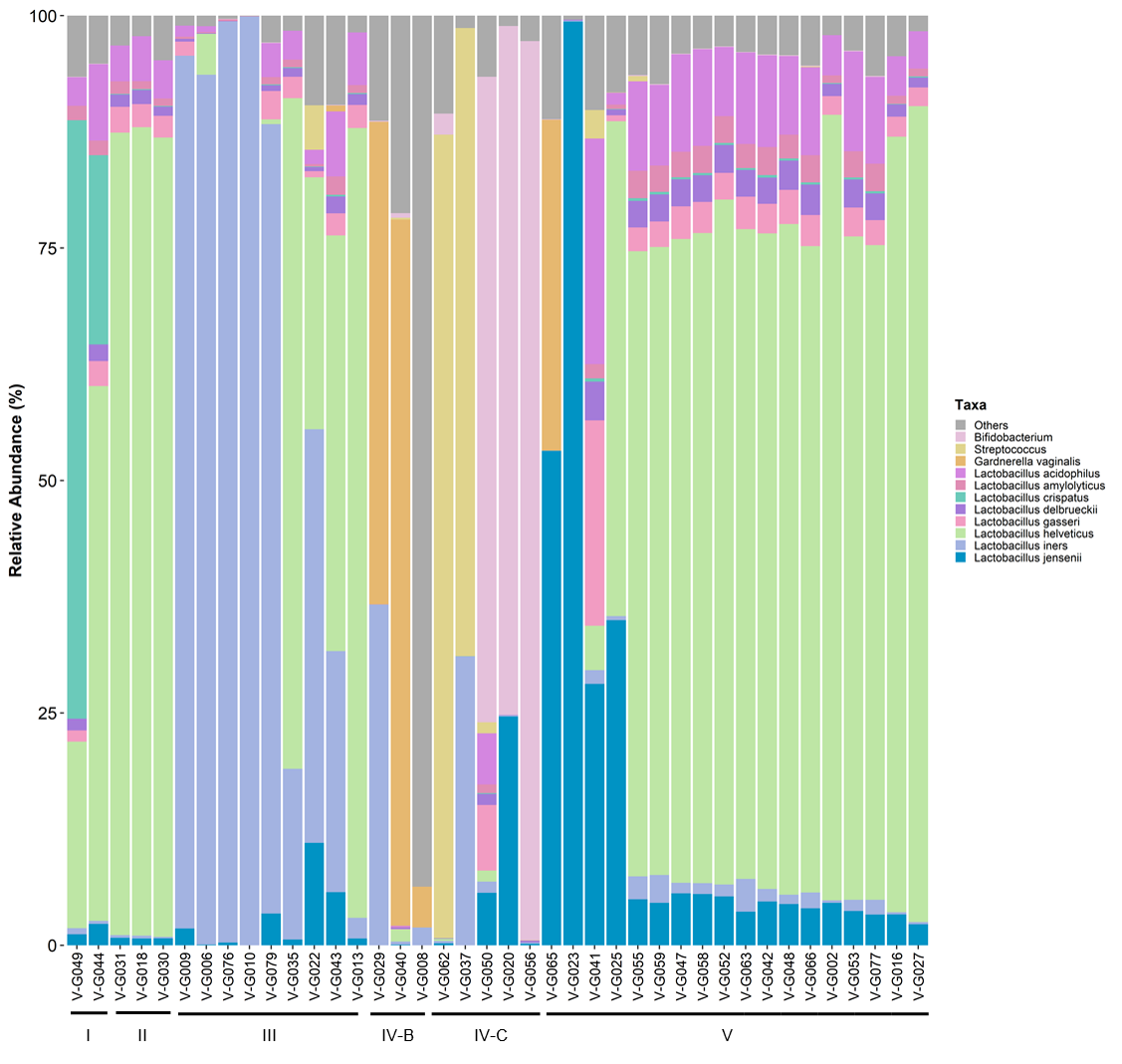

**Supplementary Figure 1.** Community state types (CSTs) of vaginal samples analysed using 16S rRNA gene sequencing in the general cohort. The microbial composition is plotted after decontamination. Bacteria with relative abundance less than 1% were grouped as “Others”.

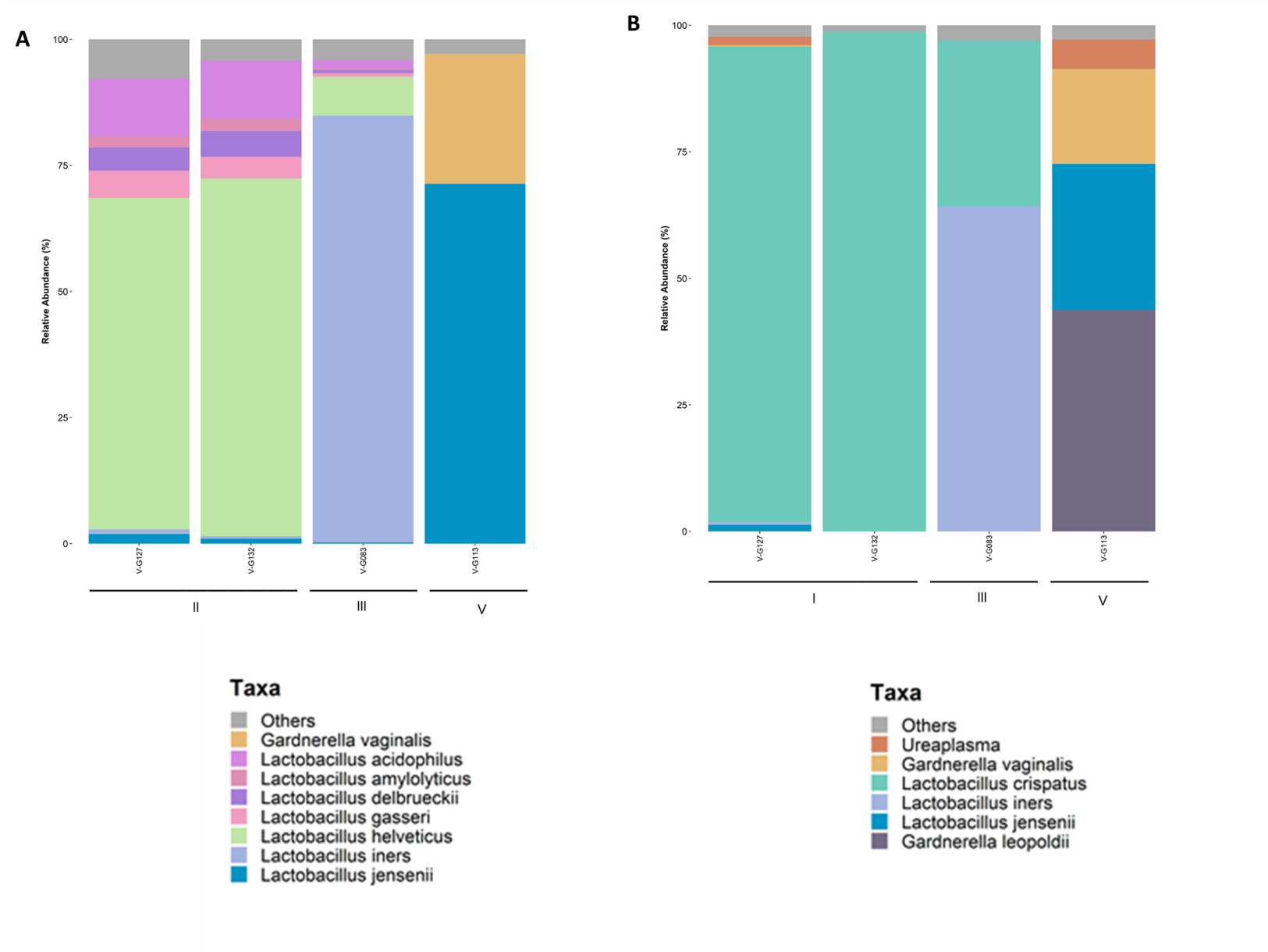

**Supplementary Figure 2.** Community state types (CSTs) of vaginal samples analysed using 16S rRNA gene sequencing (A) and meta-transcriptomics (B) in the validation cohort. The microbial composition is plotted after decontamination. Bacteria with relative abundance less than 1% were grouped as “Others”.

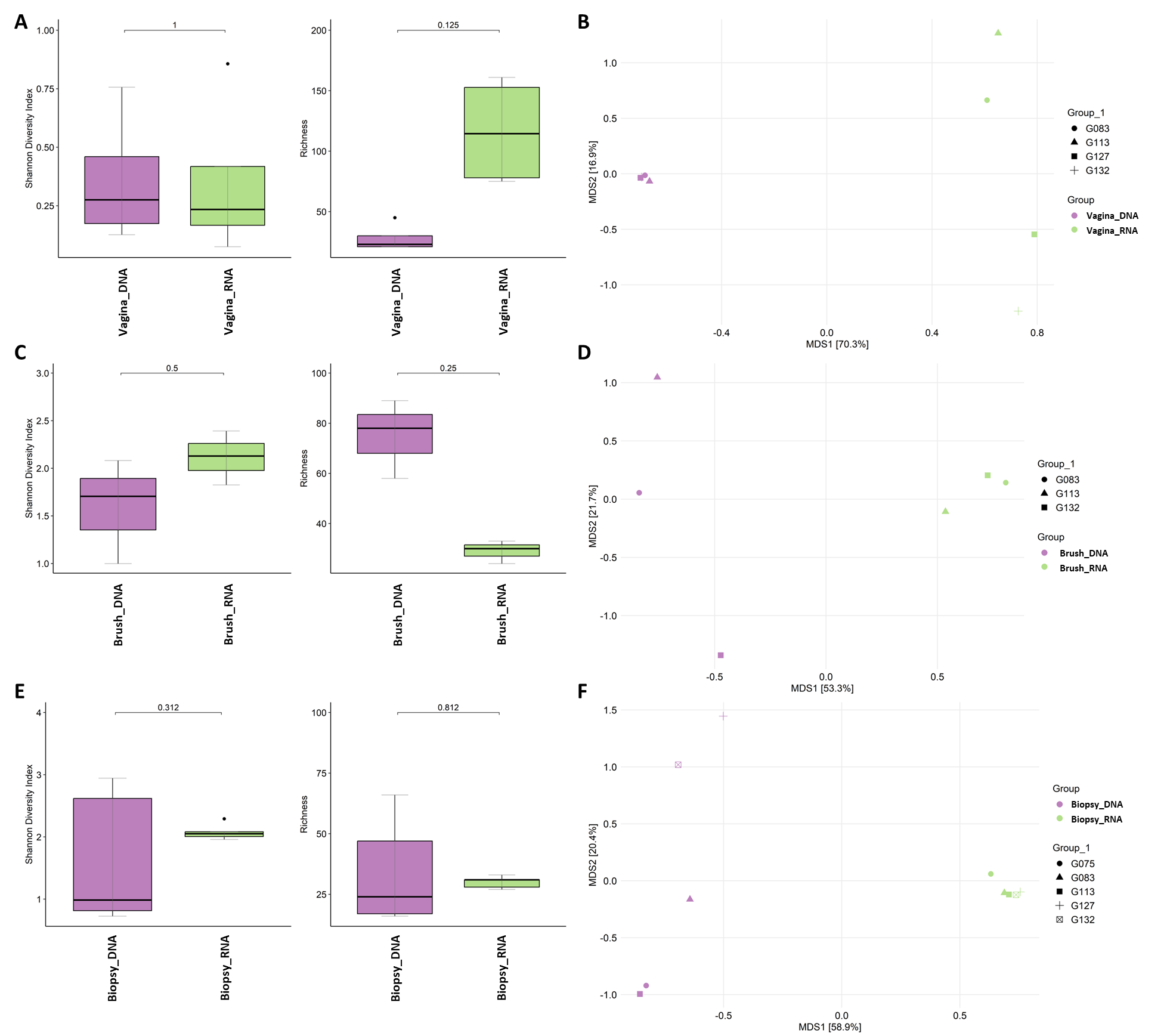

**Supplementary Figure 3.** Microbial alpha and beta-diversity measures in vaginal, endometrial brush and endometrial biopsies (DNA and RNA) samples in the validation cohort. (A, C, E) Shannon index and observed richness. (B, D, F) Principal coordinates analysis (PCoA) based on the Bray–Curtis dissimilarity (Adonis PERMANOVA, all R^2^ > 0.5, all p-values > 0.05).
